# PC_sim: An integrated measure of protein sequence and structure similarity for improved alignments and evolutionary inference

**DOI:** 10.1101/2023.01.22.525078

**Authors:** Oscar Piette, David Abia, Ugo Bastolla

## Abstract

**Motivation:** Evolutionary inferences depend crucially on the quality of multiple sequence alignments (MSA), which is problematic for distantly related proteins. Since protein structure is more conserved than protein sequence, it seems natural to use structure alignments for distant homologs. However, structure alignments may not be suitable for inferring evolutionary relationships at the sequence level.

**Results:** Here we investigate the mutual relationships between four protein similarity measures that depend on sequence and structure (fraction of aligned residues, sequence similarity, fraction of superimposed backbones and contact overlap) and the corresponding alignments. Changes in protein sequences and structures are intimately correlated, but our results suggest that no individual measure can provide a complete and unbiased picture of changes in protein sequences and structure. Therefore, we propose a new hybrid measure of protein sequence and structure similarity based on Principal Components (PC_sim). Starting from an MSA, we obtain modified pairwise alignments (PA) based on PC_sim, and from them we construct a new MSA based on the maximal cliques of the PA graph. These alignments yield larger protein similarities and agree better with the Balibase “reference” MSA and with consensus MSA than alignments that target individual similarity measures. Moreover, PC_sim is associated with a divergence measure that correlates strongest with divergences obtained from individual similarities, which suggests that it can infer more accurate evolutionary divergences for the reconstruction of phylogenetic trees with distance methods.

**Availability:** https://github.com/ugobas/Evol_div

## 1 Introduction

The study of protein evolution relies heavily on the quality of multiple sequence alignments (MSA). However, it is known that distant alignments have low accuracy with consequent errors in evolutionary inference [1–4], which partly explains why current phylogenetic methods show poor performance when applied to distantly related proteins [2]. Many MSA programs have been developed in the past years but, when they are applied to genomic scale datasets, different programs tend to produce qualitatively different conclusions [5], so that some scholars have even advocated for the need of alignment-free approaches [6]. These problems are particularly strong at the superfamily level, which are the most distant groups of proteins for which a common ancestry can be inferred and contain proteins of known structure that diversified their biological functions through long evolutionary histories [7, 8].

The importance of alignments goes beyond evolutionary studies, as many bioinformatics methods and techniques rely on them. In particular, alignment quality has a strong influence on protein structure prediction both through homology modelling [9] and through correlated substitutions [10–12], prediction of protein function [13] and molecular interactions [14]. It is thus important to improve the current multiple alignment algorithms.

Several approaches attempted to integrate structural information to improve MSA, using additional information such as for instance predicted secondary structure [15, 18] or the statistical properties of gaps in structurally aligned proteins [16, 17]. These approaches are based on the observation that protein structure is more conserved than protein sequence [19–21], so that structure similarity may still yield valuable information when sequence divergence is close to saturation. Consistently, it was found that structure-based alignments tend to be more accurate on benchmark databases, in particular for distantly related proteins and for buried residues; nevertheless, methods that combine sequence and structure information in general do not outperform structure-based methods [22].

## 2 Approach

Here we consider diverse sequence and structure similarity measures. Each of them captures correlated but different aspects of protein evolution. Therefore, we derive the hybrid sequence and structure similarity measure PC_sim that captures all of them and we show that we can use it for improving an input MSA when structure information is available.

Sequence and structure divergence provide consistent evolutionary information since they are strongly correlated [23]. However, natural selection acts with different strength on sequence and structure. The rate of structure divergence tends to be slower than sequence divergence when measured in comparable units (see Methods), in particular for proteins that conserve the molecular function [21], which suggests that natural selection constrains protein structure more strongly than sequence since mutations that conserve the protein structure but may affect other properties such as the folding stability or the metabolic cost of the protein are more frequently fixated than mutations that change the structure. Conversely, proteins that change molecular function tend to evolve faster, in particular for structure divergence, which is consistent with the idea that protein structure change is a target of positive selection [24].

There are several ways of measuring protein structure similarity and divergence, and we distinguish two main types. (1) Some similarities, such as the fraction of spatially superimposed residues, are computed after spatial superimposition, which depends on the optimal rotation matrix. A commonly applied criterion for determining this optimal rotation consists in numerically maximizing the template-model score [25] (TM, see Methods) that superimposes pairs of residues that are closer than expected by chance. The average protein coordinates in the native state allow predicting native dynamical fluctuations in reasonable agreement with experiments through the structure based Elastic network model (ENM) [26, 27]. These predicted fluctuations correlate with observed large-scale functional motions [28]. Therefore, we may expect that proteins with low TM score present very different native dynamics, as predicted through their ENM.

(2) The fraction of shared inter-residue contacts (contact overlap, CO, see Methods) is a structure similarity measures that does not require any rotation. These contacts allow estimating the folding stability of the protein through simple contact-based models [29], and we may expect that the CO correlates with the similarity of folding free energies. It is thought that protein dynamics is a target of selection more relevant than protein stability that evolves almost neutrally [30]. Consistently with this expectation, we and coworkers observed that the TM score decays more slowly than the CO in protein evolution and it is subject to stronger accelerations upon function change [24], which is consistent with the above idea that protein dynamics is subject to stronger evolutionary pressure, both negative and positive, than protein stability.

Given the intricate sequence-structure-function relationship that characterizes proteins and the fact that different similarity measures capture only part of it, we studied the correlations between these different measures. Different similarity measures give consistent information, since they are all strongly correlated between themselves. We found here that their main Principal Component (PC) represents more than 75% of the total variance, depending on the type of alignment.

We propose that the main PC of protein similarity measures (PC_sim) yields a convenient description of the similarity between proteins that integrates sequence conservation, conservation of the local coordinates (related with native dynamics), and conservation of the contact matrix (related with folding stability). We modified the MSAs obtained from popular MSA programs by targeting PC_sim as well as other structure similarity measures (TM score and CO). We found that the pairwise alignments (PA) that target PC_sim yield the highest or second highest value of all similarity scores, both for structure and sequence, and that they arguably provide better performances than alignments that target other measures. Subsequently, from the PC_sim-modified PAs we derived an MSA by determining the maximal cliques of the PA graph. We show that this modified MSA has larger similarity scores and larger similarity with reference alignments of the Balibase database [42] than the original MSA from which it is derived.

Finally, the protein similarity measures can be transformed into inferred evolutionary divergences in the same way as the Tajima-Nei divergence can be derived from sequence identity [32]. These inferred divergences may be adopted to build the guide tree for progressive multiple alignments, which has a strong influence on the final MSA and bias the phylogenetic relationships inferred from the MSA through Maximum Likelihood methods [3, 4]. We found that the divergence measure obtained from PC_sim provides the highest correlation with all other divergence measures, which suggests that it may provide a more robust inference of divergence time and guide tree.

## 3 Methods

### Alignment algorithms

We generated multiple sequence alignments (MSA) with 4 commonly used MSA programs: Clustal-Omega [35], MAFFT [34], MUSCLE [36] and T-coffee [37] and MStA with the program Mammoth-mult [38]. In all cases we used default parameters and we built the MSA using the Ebi-tool API [33].

### Protein similarity measures

For each pair of proteins with known structure, we computed the following global similarity measures, either in sequence or in structure:

- **Fraction aligned (ali)**: Fraction of positions that are considered homologs (no gaps) with respect to the maximum length of the two proteins. This normalization penalizes insertions and deletions, which can produce functional or structural changes, although different length might also come from different crystallization constructs.
- **Sequence identity (SI)**: Fraction of aligned positions that share the same amino acid (note that indels are not scored by SI).
- **TM score (TM)**: Fraction of spatially superimposed aligned positions. While the root mean square deviation (RMSD) is a good measure of structure divergence for fixed number of superimposed positions, it cannot be used when this number is variable, since there is a trade-off between the length of the superimposition and the RMSD. To address this problem, Zhang and Skolnick introduced the template-model (TM) score [25], defined as

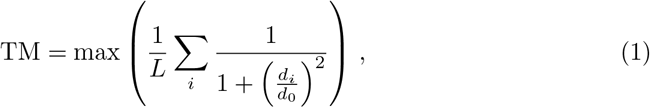

where *L* is the number of aligned positions, *d_i_* is the distance between the two alpha carbons aligned at column *i* after optimal rotation, *d*_0_ = 1.24(*L* — 15)^1/3^ — 1.8 is the L-dependent distance expected for structurally unrelated positions, and the optimal rotation matrix is determined self-consistently by iteratively maximizing the TM score.
- **Contact overlap (CO)**: Fraction of shared contacts between two aligned protein structures. Different from the TM score, the CO does not depend on any rotation matrix, and it is defined as

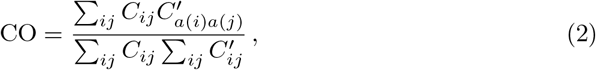

where *C_ij_* and 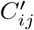 are the binary contact matrices of the two protein structures defined as 1 if any pair of heavy atoms of residues *i* and *j* are closer than 4.5Åand 0 otherwise, *a*(*i*), *a*(*j*) are the residues of the second structure aligned to residues *i*, *j* of the first one (excluding gaps). The CO is normalized so that its maximum value is 1.
- **PC similarity (PC)**. The PC is the weighted combination of the four similarity measures described above. The weights were determined through the Principal Component Analysis of the four similarity measures for all the superfamilies studied in this work. Using the MAFFT alignment to compute the similarity scores, we obtained:

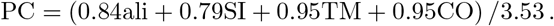

These are the weights used in this work. Other alignment programs yielded similar weights, which are reported in Supplementary Table S1.

### Evolutionary divergences

To each pairwise similarity measure we associate an evolutionary divergence that estimates the time during which the two proteins diverged. For sequence identity, we adopted the Tajima-Nei (TN) divergence [32] that represents the maximum likelihood estimate of the divergence time under the Juke-Cantor (JC) model of molecular evolution in which sites are regarded as independent, all amino acids have the same stationary frequency and all pairs of different amino acids have the same exchangeability. We adopted this estimate because it is simple and parameter-free. For the other similarity measures, we define divergences that are formally equivalent to the TN divergence. We postulate that these structural divergences can estimate the divergence time for suitably defined models of structure evolution analogous to JC, which is supported by their strong correlation with the TN divergence that we observed in previous work [21, 24].

- **TN divergence** [32] is computed from sequence identity (SI) as

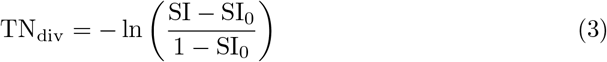

where SI_0_ = 0.05 is the sequence identity expected for unrelated sequences.
- **Contact divergence** [21], computed from the CO as

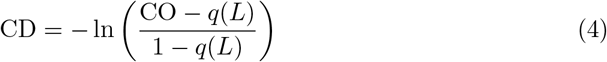

where *q*(*L*) = 0.39*L*^-0.55^ + 6.64*L*^-0.67^ is the CO expected under convergent evolution, *L* is the number of aligned residues.
- **TM divergence** computed from the TM score (TM) as

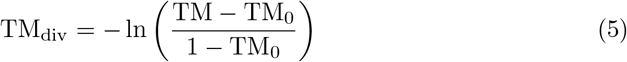

where TM_0_ = 0.167 is the TM score expected under convergent evolution [25], which is independent of *L* because the TM score is carefully normalized.
- **PC divergence**. It is a new hybrid sequence-structure divergence measure computed from the PC similarity, Eq.(3) as

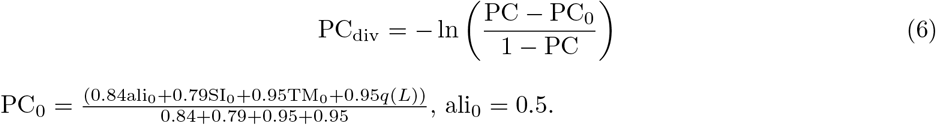

Note that the divergences can be evaluated only for similarities larger than expected under no homology (SI > SI_0_, CO > *q*(*L*), TM > TM_0_, PC > PC_0_). Structure divergences are not guaranteed to vanish for identical sequences, since the structures used to evaluate them may be related through a conformational change. To reduce this risk, we cluster the studied conformations in groups with identical sequence and we define the structure divergence between two groups as the minimum value of the divergence between their members.

### Protein superfamilies

In this work we studied four protein domain superfamilies: Globins, Ploops, NADP and Aldolases, selected as they are among the largest superfamilies in the SCOP and CATH databases [7, 8]. Proteins were parsed into globular domains in the SCOP database [7], from which we obtained the average native coordinates of the selected domains.

For each superfamily, we clustered protein structures with Contact Divergence < 2.5 as in [21], because it is not possible to obtain a Multiple structure alignment (MStA) if structural domains are too divergent. We obtained 2 clusters each for Ploops and Aldolases and one each for Globins and NADP, so that we ultimately studied 6 clusters. The distributions of the sequence and structure similarity measures for each of the four largest clusters is shown in Supplementary Fig.S1, from which one can see that sequence identity covers a broad range, with many pairs below 20% (designated as the twilight zone), most pairs between 20 and 50%, and several pairs above 50%.

Since proteins may have different conformations, we also considered proteins with identical sequences within one point mutation and clustered their structures, computing the structure similarity between clusters as the maximum across all conformations in the same cluster. The correlations and the Principal Component Analysis shown in this paper are based on the similarities between pairs of clusters. The numbers of conformations and different sequences in each cluster were the following: Aldolase C1: 38, 15; Aldolase C2: 23, 9; Globins C1: 397, 71; NADP C1: 161, 92; Ploop C1: 150, 73; Ploop C2: 45, 16.

### Modified pairwise alignments

We compute the similarity and divergence scores described above for the starting alignments as well as four new pairwise alignments (PA) modified through the use of structure information. Given the exploratory nature of the present work, we only implemented fast modified alignments that are based on the starting alignment and do not require to score gaps, which is a critical point of all alignment methods.

The first modification that we studied is the secondary structure based alignment (SS_ali). It is grounded on the idea that the sequence and the structure alignment have different aims: to infer homology and to identify structurally equivalent residues, respectively. Thus, they need not coincide everywhere. For instance, if a residue inside a secondary structure element (SSE) is deleted, the structure of the mutated residue will rearrange so to maintain the structural integrity and, in the structure alignment, the resulting gap will move to one of the two ends of the SSE. Our program detects such cases and moves the gap in the direction in which the displacement is smaller, it computes sequence identity and TM score both for the starting and the modified alignment and it selects the largest similarity. If the TM score of the modified alignment is higher, the CO is also taken from the modified alignment.

Next, we construct modified alignments that target the TM score (TM_ali), the CO (CO_ali) or the PC similarity (PC_ali) while modifying the input alignment as little as possible, through the following procedure. (1) For each residue we identify the nearest residue in the other protein as the one that maximizes the target score (CO, inter-residue distance or PCjsim), which depends on the input alignment and the optimal rotation matrix. (2) We identify as neighbors two residues that present a double match, i.e. i1 is the nearest residue of i2 and i2 is the nearest residue of i1. (3) We align neighbors that are aligned in the input alignment, obtaining frames. (4) Proceeding from left to right, we align neighbors that are intermediate between frames. We iterate this procedure, calling new neighbors using the modified alignment. In this way, we obtain modified alignments that are similar to the input alignment and increase the target score without having to specify a gap penalty parameter. This procedure provides a modified PA for each target measure.

### Multiple alignment

We then obtained an MSA from the set of modified PA through the following simple graphbased algorithm. (1) We transform the PAs into a graph with residues of the *n* proteins as nodes, whose links connect aligned residues. The maximum number of links per residue is *n*. If all PAs are consistent with an MSA, each column of the MSA corresponds to a maximal clique in the graph, i.e. a maximal set of fully interconnected residues. (2) We determine the maximal cliques of the graph for each residue *i* iteratively, exploiting the list of its neighbours limited to residues *j* > *i* in order to avoid repeated computations. The first clique is constituted by *i* and its first linked residue *l_1_*(*i*). At each step *s* we add to all previous cliques the residue *l_s_*(*i*) linked to *i*. If this residue is linked to all residues in the clique it is added to it, otherwise a new clique is created with all residues that are linked to both *i* and *l_s_*(*i*). Crucially, to reduce the computation time at each step we keep only the 100 largest cliques. When the neighbors of *i* are exhausted we store the largest cliques. The ranking is very fast because the size can only take values from 2 to *n*. For each residue we store the maximum size and the sum of the sizes of its cliques, and exclude from future computations residues for which each of them is larger than *n*/2. This precaution provides a good compromise between completeness and computational efficiency. (3) We assemble the cliques that are reciprocally consistent, i.e. they do not violate sequential order, starting from the largest one. (4) We assign the residues that are not assigned to any maximal clique to the clique most connected to them if this assignment is consistent with all other pre-existing cliques. If this is not possible, unassigned residues seed a new column. (5) We reconstruct the MSA from the set of all ordered columns (maximal cliques) and we print it for subsequent use.

### Assessment of the MSA

To assess the MSA, we downloaded the Balibase set of structure-curated multiple alignments [42]. For each MSA we only considered sequences that are associated to a protein structure in the PDB [43]. We assessed the similarity between pairs of alignments through the sum of pairs score [42] that sums the pairs of residues that are aligned in both alignments. We adopted a symmetric version of the score, normalizing the sum through the geometric mean of the sum of aligned pairs in the two alignments, which also penalizes overalignments.

### Output files

For each pair of proteins, the program Evol_div outputs for posterior analysis five similarity measures (ali, SS, TM, CO and PC) and the corresponding divergences for each of five alignments (input and modified to target SS, TM, CO and PC). The pairwise scores are printed both for all examined structures (.sim and .div) and for the clusters of structures with identical sequence (.prot.sim and .prot.div). The program also prints the MSA modified through secondary structure (ss_ali.msa) and modified by targeting the PC similarity (PC.msa).

## 4 Results

### Correlations between similarity measures

In this work we consider four protein similarity measures: fraction of aligned residues (ali), fraction of aligned residues that are identical (SI), fraction of spatially superimposed residues (TM score) and fraction of shared contacts (CO) (see Methods).

Fig.1A to D shows the similarity scores obtained with four sequence alignment programs (Clustal [35], MAFFT [34], MUSCLE [36] or T-coffee [37]) and one structure alignment program (Mammoth [38]).

**Figure 1:**
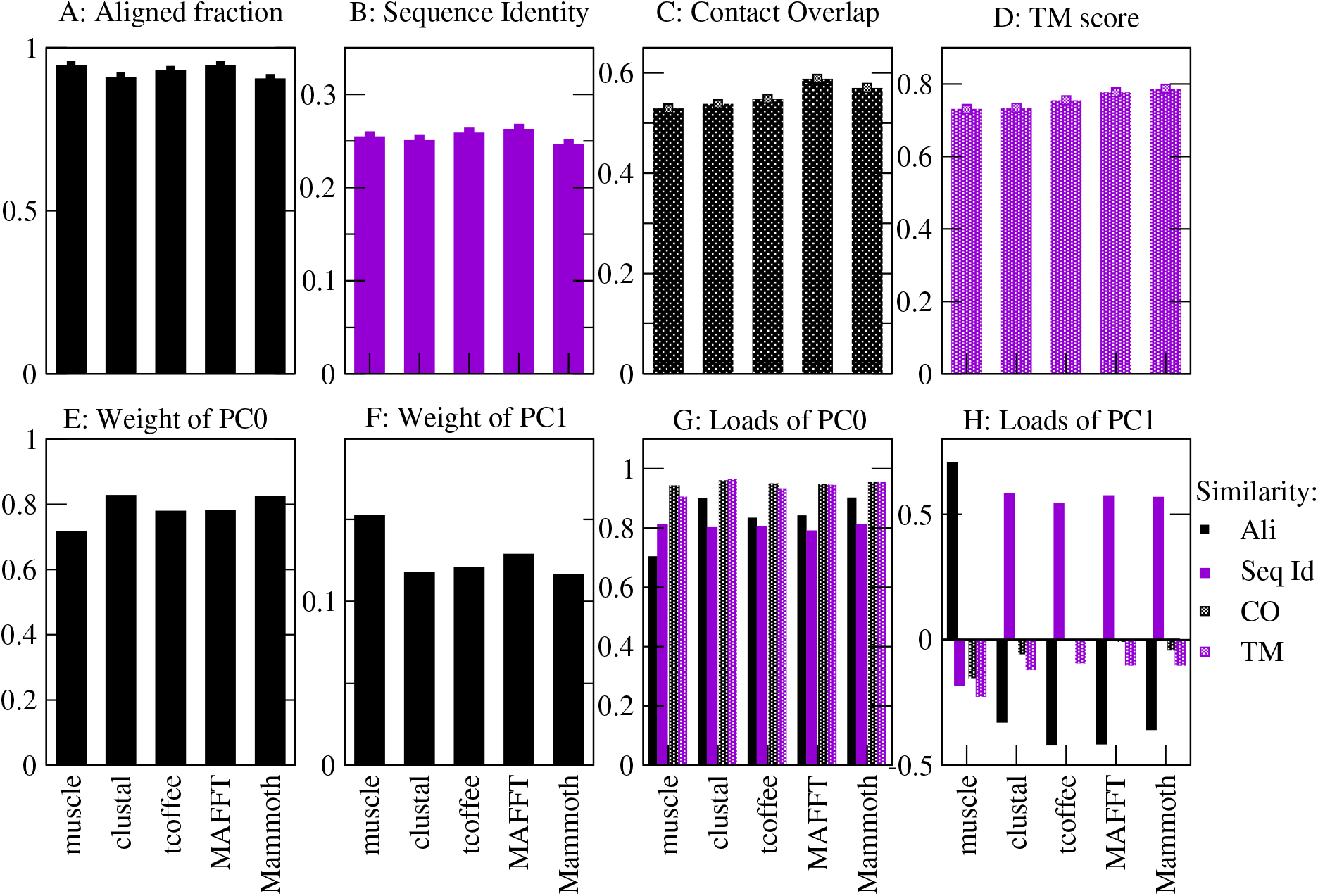
Similarity scores obtained by different alignment algorithms (A-D) and their principal components (E-H).

As expected, the sequence alignment programs attain higher sequence similarity scores and lower structure similarity scores than the structure alignment program (see Fig.1). Therefore, targeting different similarity scores with sequence alignment and structure alignment algorithms has a deep influence on the final results. Clustal and the MStA program Mammoth aligned on the average fewer residues than the other MSA programs (Fig.1A). Mammoth obtained the highest TM-scores of all programs (Fig.1D), which is not surprising since the score that it targets is related with the TM score. MAFFT achieved the second highest TM score and the highest CO, even higher than the MStA program Mammoth (Fig.1C). Due to its good performances with structural scores, sequence identity, and fraction of aligned residues (Fig.1B), in the following we present results for the MSA computed by MAFFT and the MStA computed by Mammoth if not otherwise stated.

As expected, all similarity measures are strongly correlated: two proteins that are similar because their alignment presents few gaps also present large sequence identity, large number of spatially superimposed residues and large fraction of shared contacts. Principal component (PC) analysis shows that the main PC, which we call PC0, accounts for more than three quarters of the total variance (Fig. 1E). All examined alignment programs yield similar values of the weight of PC0, with the MStA program Mammoth yielding the largest value. All similarity measures contribute positively to PC0 (Fig. 1G), and the measures that contribute most are the two structure similarity measures, which are strongly correlated between each other, while SI contributes least, except for Muscle. We can interpret PC0 as an integrated measure of evolutionary relatedness, since it is large for pairs of proteins that are strongly related under the point of view of both sequence and structure. Therefore, we adopted PC0 as a new similarity measure, PC_sim, which integrates sequence and structure conservation.

The PC loads are remarkably robust to the five used program: Their range is 0.79 — 0.81 (SI), 0.94 — 0.96 (CO), 0.91 — 0.97 (TM), with largest variation 0.71 — 0.90 for the load of the aligned fraction (see Supplementary Table S1). The loads determined with only one superfamily are very similar to those obtained with the full data set (see Supplementary Fig. S2). One might expect that the loads vary with the sequence identity, but the subsets of very distantly related pairs (SI< 0.2) and intermediate ones (0.2 <SI< 0.5) provide essentially the same loads, whereas for the small number of closely related pairs (SI> 0.5, 5% of pairs) the SI has a small load (see Supplementary Fig. S2) because it is weakly correlated with structure similarity and aligned fraction, possibly due to proteins with same sequence and different conformations. These results show that we can define the hybrid PC similarity in a robust way.

The second PC (PC1) accounts for most of the remaining variance, but its weight is much smaller (Fig. 1F). It is contributed by the ali measure and by the SI measure with opposite signs, while the structure similarities yield small contributions (Fig. 1H). This means that aligned proteins with large PC1 score have larger fraction of identical amino acids and fewer aligned residues, i.e. more gaps. This may suggest that PC1 arises from the tendency of alignment programs to overfit the sequence identity at the expense of placing gaps and slightly reducing the structure similarity. However, this interpretation is questioned by the fact that we observe PC1 also for alignments produced by the structure alignment program Mammoth that does not score sequence identity, although with the smallest weight among all examined alignment programs. An alternative interpretation is that pairs with large PC1 are domains with different size, either because they were crystallized from different constructs or because they were differently parsed in the SCOP database. Among the sequence alignment programs, the lowest weight of PC1 is attained by Clustal, followed by T-coffee, MAFFT and then MUSCLE, for which it is largest.

### 4.1 Structure-guided modified alignments

In this work, we considered four structure-guided modifications of input MSAs constructed either by the sequence alignment algorithm MAFFT [34] or by the structure alignment program Mammoth-multiple [38], which we used in previous studies and which provided good results in a recent benchmark test [22]. We obtained qualitatively similar results with both programs, and with other sequence alignment programs as well. Three modified alignments target structure similarity scores: TM_ali targets the TM score, CO_ali targets the CO, and PC_ali targets the hybrid sequence and structure similarity score PC_sim.

Figure 2 shows the differences in 5 similarity measures (ali, SI, TM, CO and PC_sim) between PC_ali and the input alignment and other three modified alignments, averaged over all pairs of alignments. One can see that the targeted score is always highest in the alignment that targets it compared to other alignments. However, compared with PC_ali, this improvement happens at the expense of all other scores. PC_ali obtains the first or second highest score for all similarity measures, both sequence and structure, except a small decrease in the aligned fraction when the input alignment is MAFFT. Globally, PC_ali was the modification with the highest average improvement of the similarity measures.

**Figure 2:**
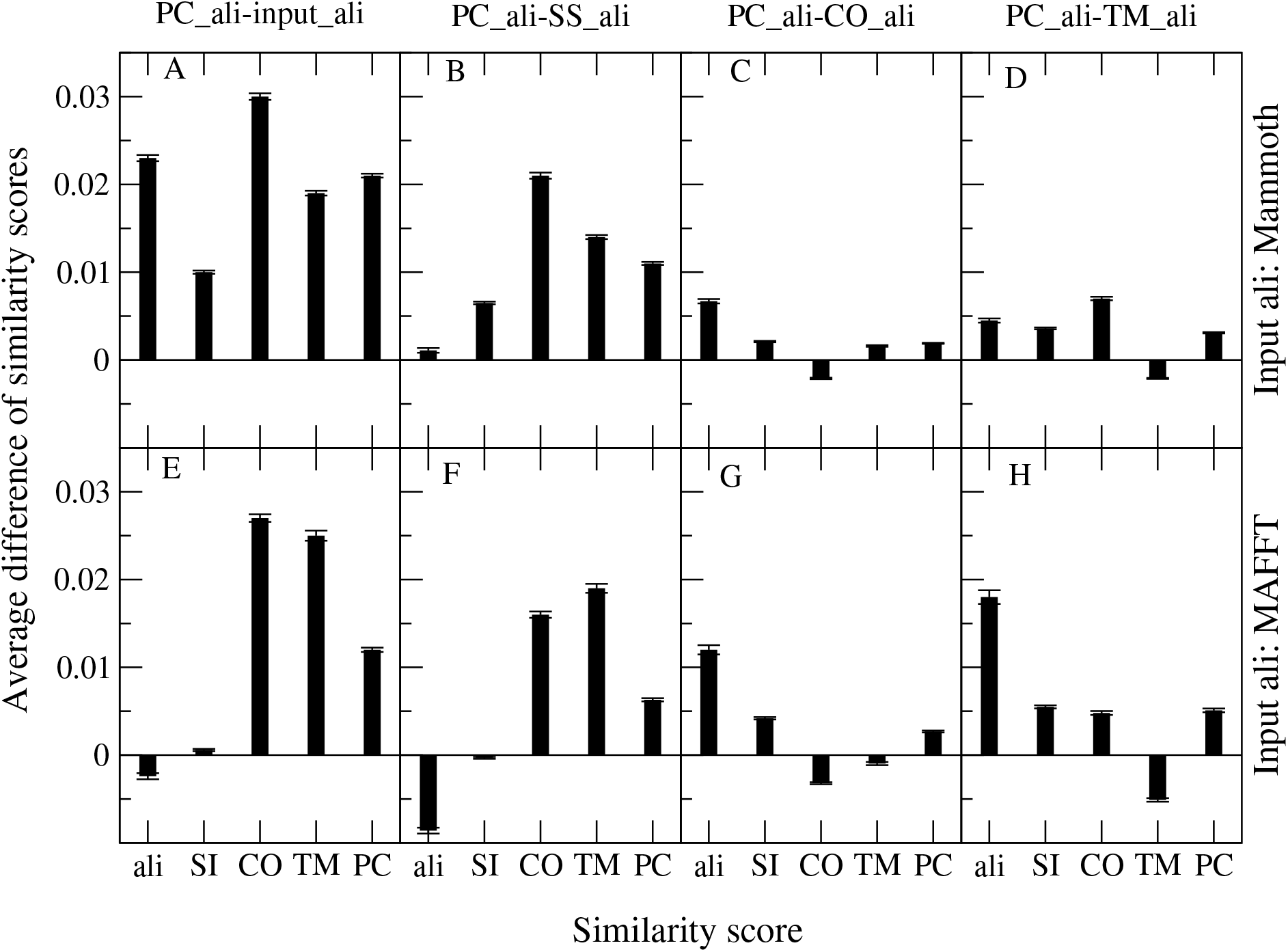
Average difference between the similarity scores obtained through the PC_sim modified alignment (PC_ali) and the input alignment (A,E) and three additional modified alignments (B,F: SS_ali; C,G: TM_ali; D,H: CO_ali) with respect to fraction of aligned residues (ali), identical amino acids (SI), spatially superimposed residues (TM), shared contacts (CO), and the hybrid PC_sim that integrates all of them (PC). The upper plots show the case in which the starting alignment is Mammoth, in the lower plots it is MAFFT. The error bars indicate the standard error of the mean.

Note that, when using as input the MStA obtained through Mammoth, the PC_ali correction increase all the sequence and structure similarities, which suggests that the alignment quality overall improves. PC_ali obtains the second highest structural score TM and CO, the highest sequence score SI and aligned fraction and, not surprisingly, the highest hybrid score PC_sim. Of note, a recent paper reported that hybrid sequence-structure alignment methods performed worse than the Mammoth program [22], which is not the case for the approach based on PC_ali.

The other modification (SS_ali) moves the gaps contained inside any secondary structure element (SSE) towards the closest end of the SSE. This is motivated by the idea that gaps inside a SSE might happen in evolution, so that the sequence alignment correctly infers homology, but the structure reorganizes so that the structural correspondence is different from the one dictated by the sequence alignment, i.e. sequence and structure alignment do not need to coincide. Accordingly, when we compute the average similarity scores we consider the higher between the SI score of the starting and the modified alignment, and the higher between the two TM scores. Not surprisingly, this procedure increases all similarity scores in Fig.2. To test our interpretation, we consider the effect of SS_ali on the similarity scores without selecting the higher score. If SS_ali is capturing gaps inside SSE, we expect that the SI tends to decrease when the structure similarity scores TM and CO increase. Nevertheless, contrary to our interpretation, we found that in most cases SS_ali decreases the similarity scores SI, TM and CO, both with respect of the sequence aligner MAFFT and with respect to the structure aligner Mammoth, i.e. modifications that improve the structure similarity are less frequent (Supplementary Fig.S3A). Moreover, sequence identity and structure similarity tend to increase or decrease together (see Supplementary Fig.S3B), which suggests that SS_ali is either correcting alignment errors through the use of secondary structure information or it is creating mistakes, instead of dealing with genuine cases of indels inside SSE that motivated it.

The best results are obtained with MAFFT as input alignment, which achieves the highest PC_sim followed by Mammoth (see Supplementary Fig. S4). Interestingly, the scores obtained with PC_ali are more robust with respect to changes of the input alignment than the scores obtained with the input alignment itself. In particular, using MAFFT or Mammoth as input alignment does not have a significant influence on the score PC_sim (see Supplementary Fig. S4).

### Generation and assessment of the MSA

As explained in the Methods section, we transform the PC-modified pairwise alignments into a graph and we determine its maximal cliques, from which we generate an MSA. We assess these PC and clique-derived MSAs by comparing them with the structure-curated MSAs of the Balibase data set [42], retaining only sequences with available structure in the PDB.

As a preliminary step, we assessed the quality of the Balibase alignments against the structure alignments obtained through Mammoth. Not surprisingly, the Mammoth alignments have significantly higher structural scores in terms of TM-score (mean difference Δ = 0.013, Standard Error of Mean 0.005) and, not significantly, in terms of contact overlap (Δ = 0.0056, SEM= 0.005), while the Balibase alignments have significantly higher fraction of aligned residues (Δ = 0.023, SEM= 0.006) and sequence identity (Δ = 0.017, SEM= 0.002), resulting in not significantly different PC scores (Δ = 0.0046, SEM= 0.004), see Fig.3M and N. As for all other alignments, the structural scores of the Balibase alignments can be improved through our algorithm (Fig.3S). Furthermore, we found bugs in Balibase sequences, since they omit residues whose index in the PDB presents an insertion code (this relatively frequent situation affects 8% of the PDB sequences in Balibase). Rarely, Balibase sequences include more than one chain when the order of the chains in the PDB file is distinct from the alphabetic order. These discrepancies forced us to realign the Balibase alignments with the modified alignments produced by our program, which are based on the PDB sequences.

**Figure 3:**
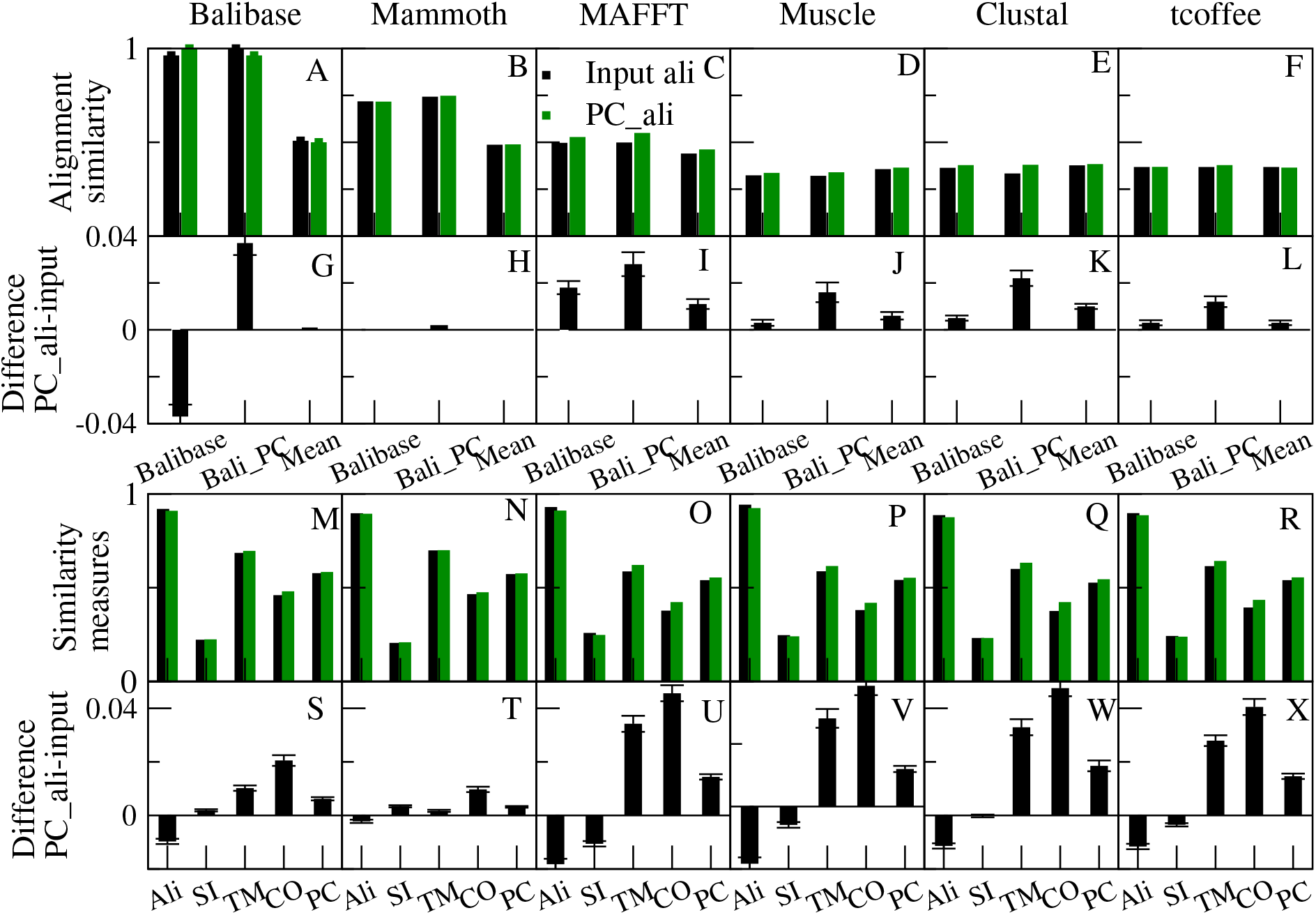
Comparison between the input MSA and the PC-modified, clique-derived MSA of 146 protein sets of the Balibase database with at least 2 protein structures. Plots A-F: absolute values and G-L: differences between alignment comparison scores (first column: Balibase reference; 2nd column: PC-modified Balibase; 3rd column: mean of all MSA comparisons). One can see that the PC correction improves the comparison with other alignments, except for Balibase and Mammoth. Plots M-R: absolute value and S-X: differences of the protein similarity scores. One can see that the PC correction improves the structural scores TM and CO for all input MSAs, at the cost of decreasing the aligned fraction. The sequence identity decreases for the sequence aligners MAFFT, Muscle and Tcoffee, but it improves for the structure aligner Mammoth and for Balibase. The overall balance, as assessed through PC_sim, is always positive. The error bars denote the standard error of the mean and allow to visually assess the significance.

Therefore, in addition to adopting the Balibase alignment as the reference alignment, we also adopted as reference the PC-modified Balibase alignment that has higher structural scores, and we adopted a consensus assessment based on comparing all alignments against all (the six input alignments Balibase, Mammoth, MAFFT, muscle, Clustal and Tcoffee, and their modified versions, omitting the comparison between each alignment and its modified version). All three comparisons show that the modified alignments have significantly higher similarity than the original alignments, with the exception of the modified Mammoth alignment for which the difference is not significant (Fig.3G-L). Moreover, for all input alignments including Balibase and Mammoth, the structural scores of the modified alignments improve at the price of producing shorter alignments (Fig.3G-L). We think that the structural scores provide a less biased assessment of the alignment quality than choosing a golden reference or a consensus, which may be biased if most of the alignments are biased in a similar direction.

### Divergence measures

The results presented above suggest the existence of compensatory changes, particularly strong for closely related protein pairs, that make difficult to disentangle the evolutionary history of a protein superfamily in terms of only one divergence measure (e.g., at the level of amino acid identity, or 3D superimposition, or contact divergence). These observations support our proposal to adopt a hybrid measure that integrates various aspects of protein sequence and structure similarity, such as the PC_sim measure presented in this paper. We now assess whether the new similarity measure can improve our ability to infer protein divergence.

From the comparison of the aligned sequences we can infer the time past since the divergence of the two proteins using simple substitution models. This inference can be expressed by simple measures such as the Tajima-Nei (TN) divergence [32]. The estimated divergence is often used to construct a guide tree for guiding the processive multiple alignment algorithm, therefore its accuracy has an important influence on the final results.

Adopting the TN formula, we can estimate divergence times using other structure divergence as well. Although the analogy is only formal, we expect that these measures may be also derived from simple probabilistic models of protein structure evolution. Since all divergence measures aim at inferring the same quantity, we can estimate their quality by assessing the strength of their reciprocal correlations.

We compute these correlations through a linear model with an offset, *D*2 = *aD*1 + *b*. In principle the offset *b* should vanish, because the divergence *D*2 should vanish for *D*1 = 0. However, protein structures may differ even for identical sequences due to the presence of conformational changes or the influence of different experimental condition on the structure determination, so that in practice the offset *b* is never zero for structural divergence, even if it is minimized by our approach to consider the maximum structural similarity over all conformations of the same protein.

For every divergence measure we compute the average correlation coefficient with the other divergences, which we present in Fig.4. High average correlation means that the divergence measure can be used to reliably estimate the other measures. The measure with highest correlation allows to reliably predict the other measures and it is expected to provide the most reliable inference of the divergence time.

**Figure 4:**
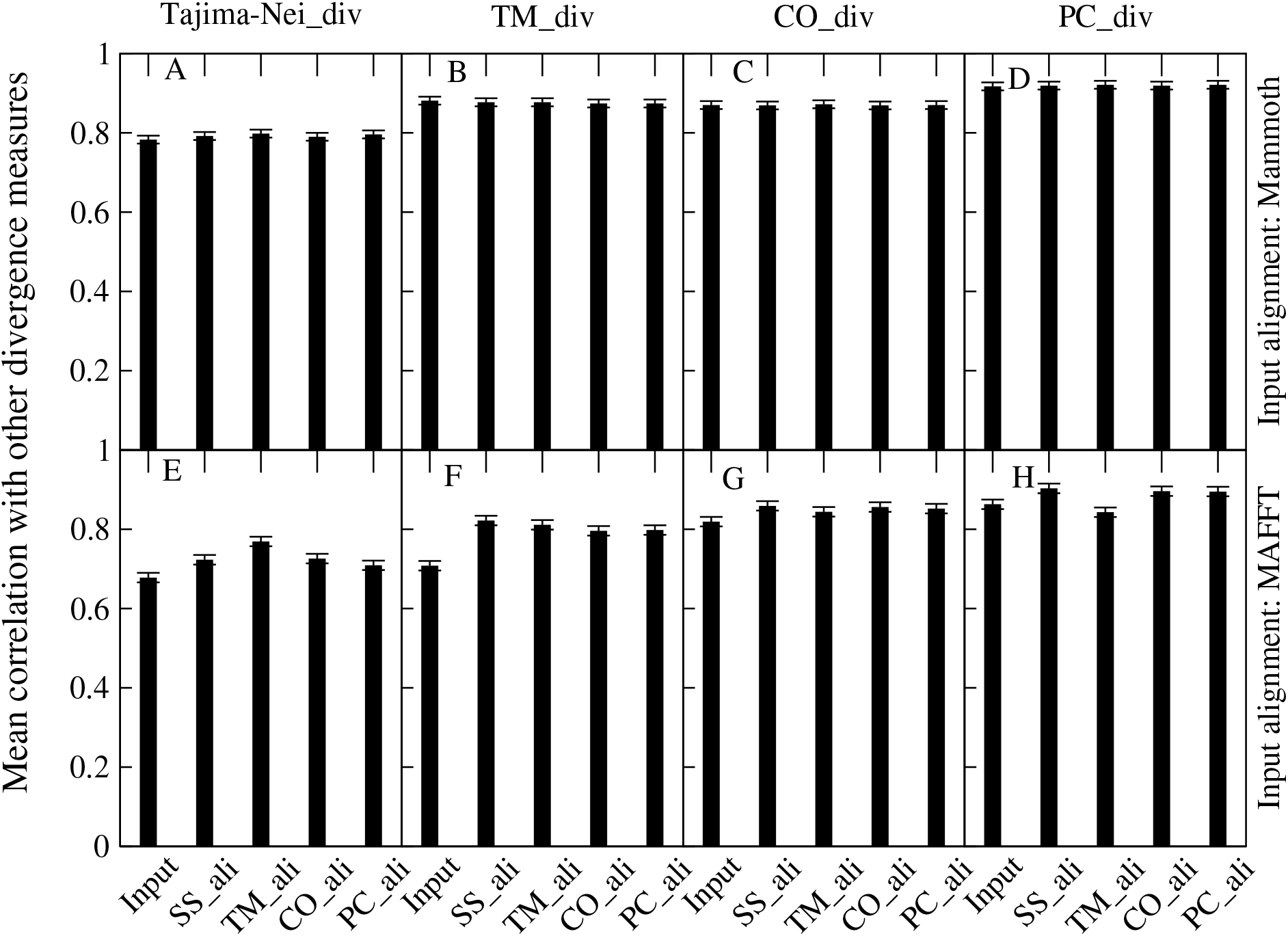
Correlation coefficients between different divergence measures for different types of modified alignments. In the top row the input alignment is Mammoth, in the bottom row it is Balibase. For all modified alignments and input alignments the highest correlations are attained with PC_div (D,H), and the lowest ones with the purely sequence-based Tajima-Nei divergence (A,E).

We see from Fig.4 that the sequence-based TN divergence Eq.(3) has the lowest average correlation with the other measures, followed by the divergences of the structural scores TM score Eq.(5) and contact overlap Eq.(4). The highest correlation (0.92 for alignments derived from MStA) is attained by the divergence of the hybrid sequence and structure similarity PC_sim (PC_Div), Eq.(6), which is therefore expected to provide the best inference of the divergence time among all divergence measures that we examined. The five modified alignments yield the same ranking of the divergence measures (Fig.2), with the input alignment MAFFT generally providing lower correlations than the modified alignments whereas the correlations obtained starting from the MStA program Mammoth are higher and quite robust with respect to the modified alignment.

## 5 Discussion

Alignment methods optimize the global similarity between aligned positions, defined either in terms of sequence or in terms of structure. Structure similarity can be measured either in terms of atomic coordinates superimposed through an optimal rotation (as in the TM-align [40] or Mammoth [38] programs) or in terms of inter-residue contacts that are independent of rotations (as in the Dali program [41]).

Here we addressed the influence of the similarity score that is targeted by an alignment program. Different similarity scores tend to be correlated, as expected if similarity is inversely correlated with evolutionary divergence (see for instance [21]). This suggests that different criteria tend to identify aligned positions in a consistent way. However, the correlations between similarity measures are not perfect, and they may produce systematically different evolutionary inferences, since the adopted similarity measure has a strong influence on the resulting inferred homology. Sequence alignments and structure alignments tend to present important differences, in particular for distantly related proteins, as well as structure alignment programs based on optimal rotation matrices or based on contacts.

In order to get more insight on the agreement and disagreement between different measures of protein similarity and evolutionary change, we performed a large scale analysis of the correlations between conservation and changes of different properties over four large protein superfamilies, i.e. homologous proteins with known structures that have diverged in sequence, structure and function throughout a long evolutionary story [7, 8]: Globins, Aldolases, P-loop and NADPH.

First of all, we confirmed that the global conservation scores of different properties are correlated. These correlations can be exploited for constructing a new integrated similarity measure based on the main principal component of both sequence similarity and structure similarity measures, see Fig.1. We called this new hybrid score “PC_sim”.

We then constructed three new alignments that modify the starting MSA (produced either by the sequence aligner MAFFT [34] or by the structure aligner Mammoth-multiple [38]) by targeting three different similarity measures: rotation-dependent structure similarity measured by the TM score [25] Eq.(1), rotation-independent structure similarity measured by the contact overlap Eq.(2), and the hybrid sequence and structure similarity measure PC_sim. Our algorithm, described in the Methods section, is based on the identification of structural “neighbors” through double best match, it does not require to determine new gap parameters, and it produces modified pairwise alignments that minimally modify the input alignment.

Our analysis supports the idea that different properties tend to give consistent information, but not exactly interchangeable. In fact, compensatory mechanisms may reduce the correlation between similarity measures: for instance, contact conservation may be achieved through compensatory changes of the coordinates of the residues in contact. Therefore, we expect that no individual similarity measure can give an unbiased description of protein evolution, and it is useful to combine different measures, as we do with our new integrated similarity measure PCjsim, in order to exploit the synergies that exist among them. Targeting PC_sim increases not only the targeted measure, but all of the structure similarity measures that we examined, including sequence similarity.

The targeted similarity measure has a systematic influence on the similarity scores, see Fig.2. Targeting structure similarity with input MSA derived from MAFFT tends to decrease the sequence identity, except with SS_ali and with PC_ali that targets PC_sim, which considers sequence identity. The two purely structural modifications increase the TM score and the CO at the expense of sequence similarity and aligned fraction, with no effect on PC_sim for the alignments that target TM and a moderate increase of PC_sim for the alignment that targets CO. These results suggest that TM_ali may be overfitting the TM. On the other hand, the alignment PC_ali that targets PC_sim improves or maintains all similarity measures, and arguably it has the best performances.

When starting from the MStA built by Mammoth, the targeted alignments improve both the sequence similarity and the structure similarities, and PC_ali achieves the best improvement for sequence identity, aligned fraction and PC_sim and the second best improvement for the TM score and the CO, again suggesting that it outperforms the other targeted alignments. Note that that a recent study found that hybrid sequence-structure alignment methods performed worse than the Mammoth program [22], while our PC_sim-based approach largely improves upon Mammoth results. Moreover, the alignments that target PC_sim are more robust with respect to variation of the starting MSA than the starting MSA themselves, which supports their use.

Our results suggest that the hybrid sequence and structure alignment method based on optimizing PC_ali can produce high quality alignments. We plan to work in the future at developing a progressive MSA algorithm that adopts an evolutionary treatment of the indel process. In this work, we obtained graph-based MSAs as the sets of the maximal cliques of the graph of the PC-corrected pairwise alignments, and we assessed them on the Balibase structure-curated MSAs [42]. On the average, the PC correction improves the similarity of the input MSA with the Balibase MSA, with the PC-corrected Balibase MSA, and the mean over all MSAs. More importantly, it improves all structural scores and the hybrid PC_sim measures (Fig.3). This supports the use of the PC-corrected MSAs.

We also constructed the modified alignment SS_ali based on moving the gaps that occur inside secondary structure elements (SSE) at the end of these elements. We reasoned that it is possible that an indel occurs inside the SSE, and the sequence alignment that infers homology should reflect it, but the native structure arranges to preserve structural integrity so that in this case sequence alignment and structure alignment do not need to coincide. If SS_ali is accounting for these cases, we would expect that increases of the TM score tend are associated with decreases of sequence similarity. However, we observed the opposite (see Supplementary figure S3): Instances in which the TM score improves and the sequence similarity decreases are less frequent than expected by chance, while most changes tend either to increase or to decrease both sequence similarity and TM score at the same time, suggesting that they either correct mistakes in the alignment or create mistakes. This is consistent with the result that gaps tend to occur rarely in SSE [16]. The same results were obtained using input alignments based both on sequence and on structure, and they do not support the use of SS_ali, which was outperformed by the modification based on PC_sim.

Last, we tested whether PC_sim improves the inference of the evolutionary divergence time. To this aim, we adopted simple estimates of the evolutionary divergence based on protein similarity measures, formally analogous to the Tajima-Nei divergence measure obtained from protein identity and often used in evolutionary studies [32]. We previously introduced a divergence measure based on contact divergence [21] and one based on the TM score [24], finding that these divergence measures are strongly correlated with each other, as expected if they are both correlated with the evolutionary time that we aim to infer. The divergence measure that is most strongly correlated with all others may provide the most robust inference of the evolutionary time. We found that the Tajima-Nei divergence shows the weakest correlations, while purely structure divergence measures are intermediate and the hybrid sequence and structure measure PC_div, based on PC_sim, shows the strongest correlations (see Fig.4), suggesting that PC_sim is able to better infer the evolutionary divergence time and, consequently, to produce better guide trees for progressive multiple alignments. The construction of these progressive multiple alignments based on PC_sim and of the corresponding trees will be the subject of our future work.

## Funding

This work has been supported by the grant PID2019-109041GB-C22/10.13039/501100011033 of the Spanish Agency of Research (AEI). Research at the CBMSO is facilitated by the Fündación Ramón Areces.

## Notes

### Competing Interest Statement

The authors have declared no competing interest.

