## Supplementary figures for "PC_sim: An integrated measure of protein sequence and structure similarity for improved alignments and evolutionary inference"

### Supplementary Material for the paper: PC_sim: An integrated measure of protein sequence and structure similarity for improved alignments and evolutionary divergence

Oscar Piette(1), David Abia(1) and Ugo Bastolla(1,2)

(1) Centro de Biologia Molecular "Severo Ochoa"

CSIC-UAM Cantoblanco, 28049 Madrid, Spain

(2)

October 27, 2022

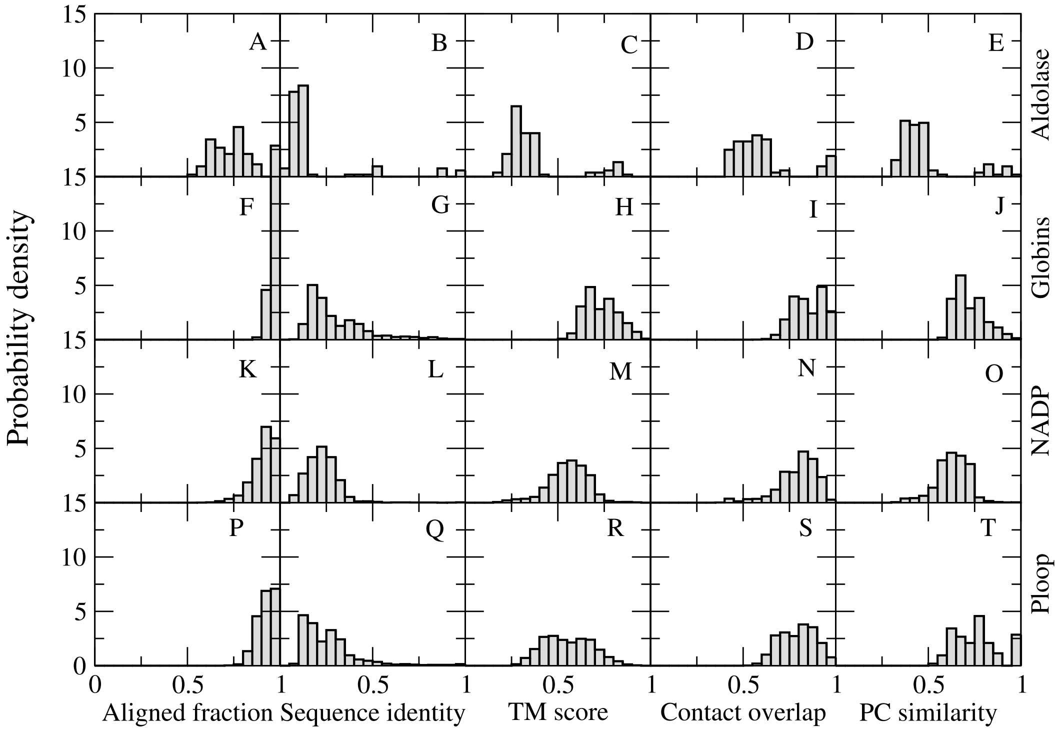


Figure 1: Distribution of the aligned fraction (1st column), Sequence identity (2nd column), TM score (3rd col), Contact overlap (4th col) and PC similarity (5th col) in the superfamilies Aldolase (110 pairs, plots A-E), Globins (2490 pairs, plots F-J), NADP (4191 pairs, plots K-O) and Ploop (2633 pairs, plots P-T). One can see that the four superfamilies span a very broad range of sequence identity.

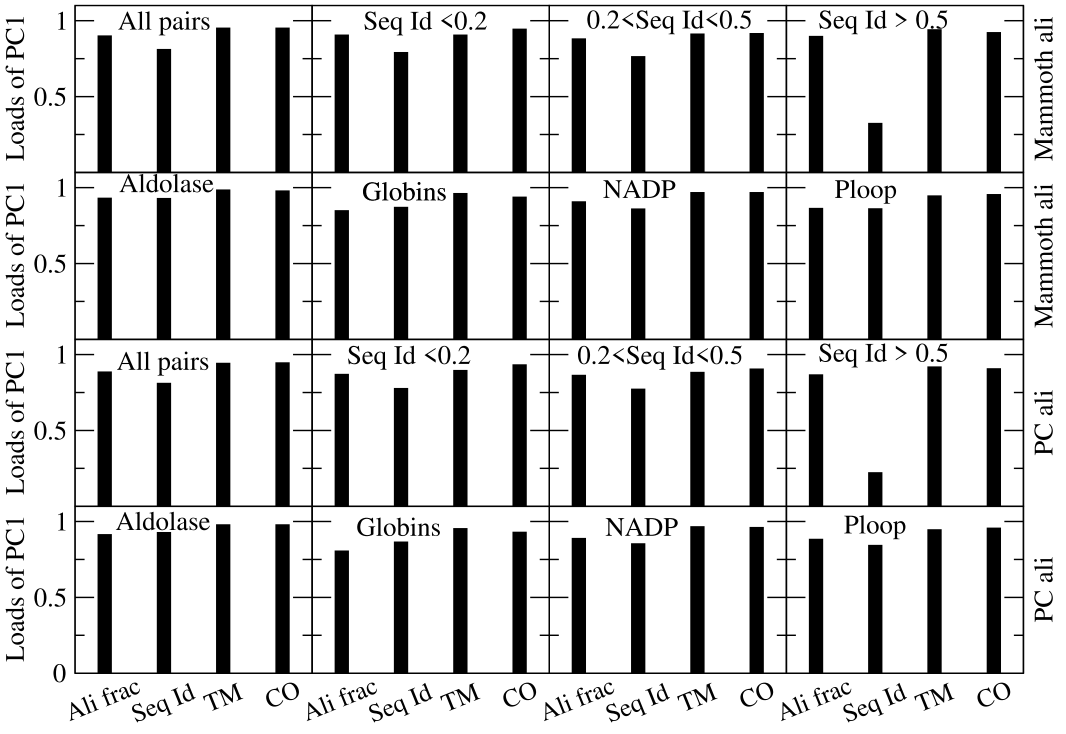


Figure 2: Loads of the first principal component of protein similarity measures (PC_sim) computed using all data (top left), pairs with low, intermediate and high sequence identity (top row) and pairs belonging to each of the four superfamilies (second row). The first two rows are obtained with the original Mammoth alignments, and the third and fourth are obtained with PC corrected Mammoth alignments. One can see that the PC loads are relatively robust, except that the contribution of sequence identity decreases at high sequence identity (5% of the pairs) due to its low correlation with aligned fraction and structure similarity measures, possibly because of the presence of conformational changes in the data set.

Table 1: Loads of the four similarity measures for the computation of the first principal component PC_sim from different input alignments.

| Program | Ali | SI | CO | TM |
|---|---|---|---|---|
| Muscle | 0.71 | 0.81 | 0.94 | 0.91 |
| Clustal | 0.90 | 0.80 | 0.96 | 0.97 |
| T-Coffee | 0.84 | 0.81 | 0.95 | 0.93 |
| MAFFT | 0.84 | 0.79 | 0.95 | 0.95 |
| Mammoth | 0.90 | 0.81 | 0.96 | 0.96 |

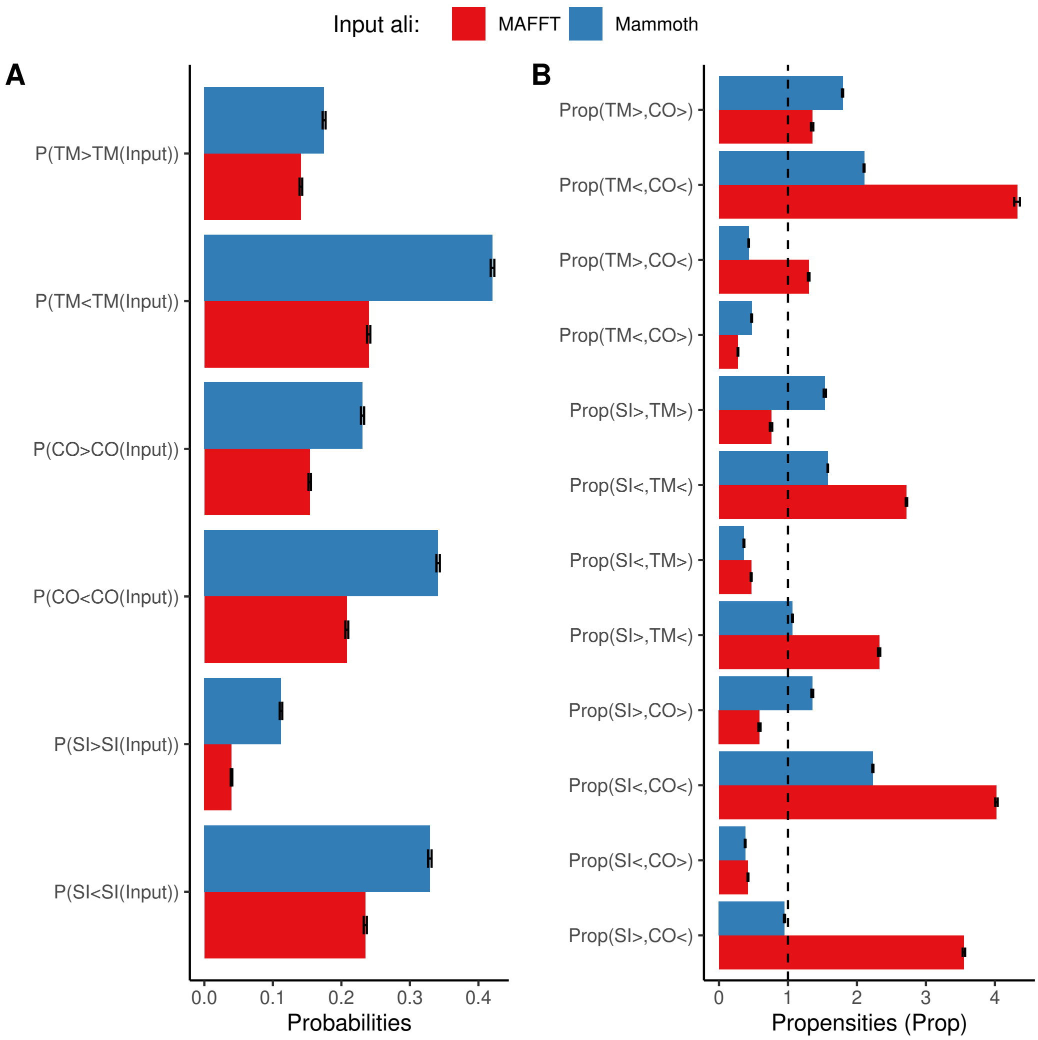


Figure 3: Performances of the SS_ali alignment modified based on secondary structure. SS_ali moves the gaps contained inside any secondary structure element (SSE) towards the closest end of the SSE, motivated by the idea that gaps inside a SSE might happen in evolution, so that the sequence alignment correctly infers homology, but the structure reorganizes so that the structural correspondence is different from the one dictated by the sequence alignment. If SS_ali is capturing such events, we expect that the SI tends to decrease when structure similarity scores TM and CO increase. Panel A represents the percentage of protein pairs for which any similarity increases or decreases with respect to the starting alignment, and it shows that in most cases SS_ali decreases the similarity scores SI, TM and CO, both with respect of the sequence aligner MAFFT and with respect to the structure aligner Mammoth. Panel B represents the propensities of all types of changes of similarity scores to occur together. It shows that increases of SI tend to be associated with increases of the structural scores TM and CO, and decreases of SI tend to be associated with decreases of the structural scores. This association is found both for starting alignments obtained with MAFFT and with Mammoth, and it suggests that SS_ali is either correcting alignment errors through the use of secondary structure information or introducing mistakes instead of dealing with genuine cases of indels inside SSE that motivated it.

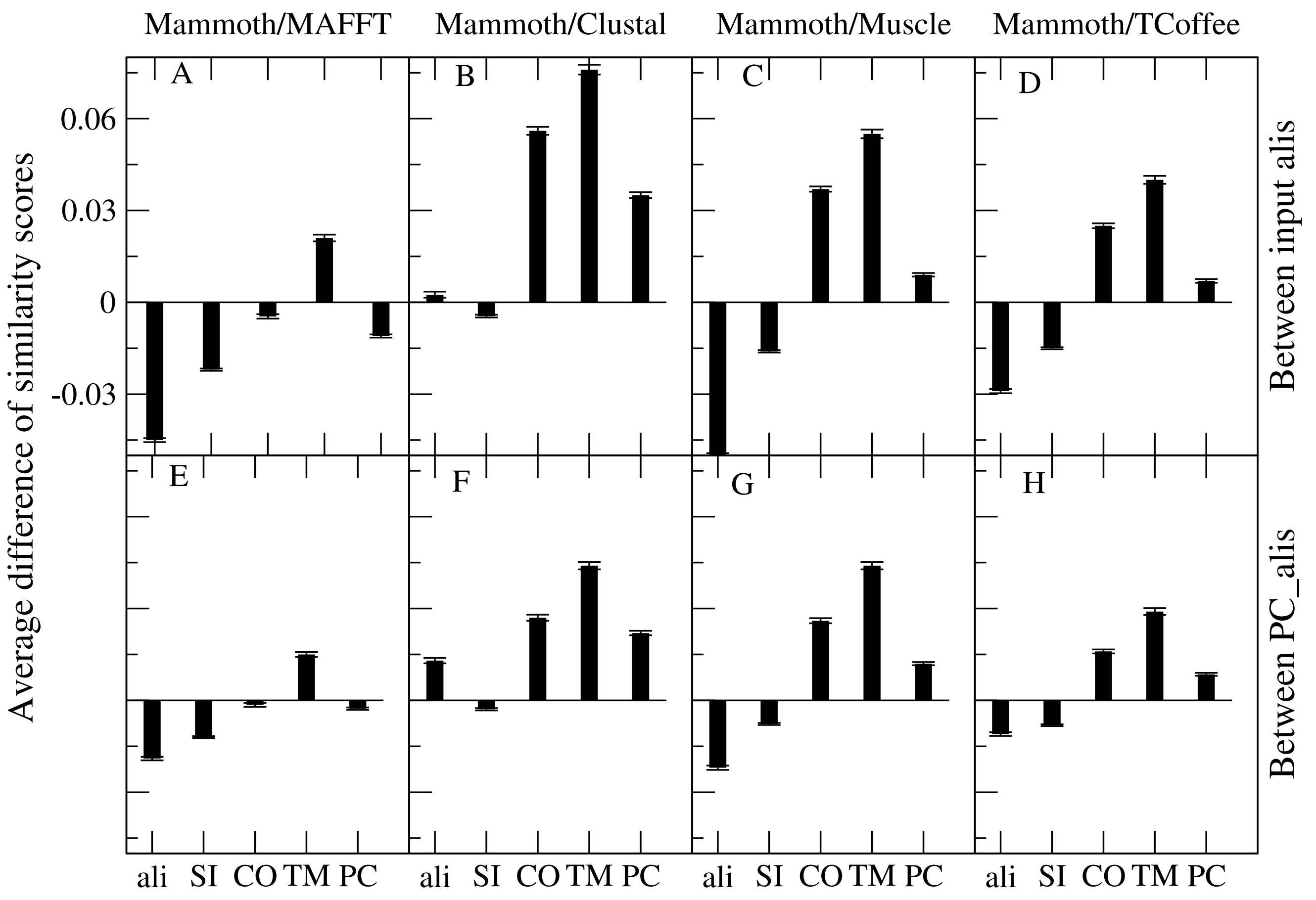


Figure 4: Influence of the input alignment on the similarity scores. Mammoth as input ali is compared to MAFFT (A,E), Clustal (B,F), Muscle (C,G) and Tcoffee (D,H). On the top line (A-D) the input alignments itself are compared, while on the bottom line (E-H) the PC_ali derived from them are compared. One can see that the PC-modified alignments vary less than the corresponding input alignments (compare the top and bottom line). Note that MAFFT provides CO scores and PC scores higher than the structure aligner Mammoth, but the difference decreases after applying the PC correction.
